# The influence of unmeasured confounding on the MR Steiger approach

**DOI:** 10.1101/2021.10.28.466267

**Authors:** Sharon M. Lutz, Kirsten Voorhies, Ann Chen Wu, John Hokanson, Stijn Vansteelandt, Christoph Lange

**Author notes:** For Correspondence: Sharon Lutz PhD, Department of Population Medicine, Harvard Medical School and Harvard Pilgrim Health Care Institute, 401 Park Drive, Suite 401 East, Boston, MA 02215.

## Abstract

The Mendelian Randomization (MR) Steiger approach is used to determine the direction of a possible causal effect between two phenotypes [1]. For two phenotypes, denoted phenotype 1 and 2, the MR Steiger approach is composed of two parts: (1) MR is performed for a set of single nucleotide polymorphisms (SNPs) that serve as instrumental variables for phenotype 1 and (2) the difference of two correlations, the correlation between the SNPs and phenotype 1 and the correlation between the SNPs and phenotype 2, is calculated. These two parts are then used to determine the direction of a possible causal effect between the two phenotypes. The original MR Steiger paper [1] shows that unmeasured confounding of the two phenotypes affects the validity of the MR Steiger approach, but does not elucidate as to how this occurs. In particular, it was argued that if the magnitude of the observational variance explained between the two phenotypes is above 0.2, the MR Steiger method may return the incorrect causal direction due to unmeasured confounding. This may initially seem surprising since unmeasured confounding does not induce spurious associations between the SNP and phenotype 2, as we demonstrate using directed acyclic graphs. In this note, we show that this is because unmeasured confounding may rescale the magnitude of a non-zero association, and thereby distort the comparison of the correlation between the SNP and phenotype 2 and the correlation between the SNP and phenotype 1. We will end with a number of cautionary remarks on the MR Steiger method, which are partly motivated by this and mentioned in the original MR Steiger paper [1].

---

Let *g* denote the SNP, *x* denote phenotype 1, *y* denote phenotype 2, and *u* denote an unmeasured confounder of the association between both phenotypes. Then, *x* and *y* can be modeled as follows in the original MR Steiger paper [1]:

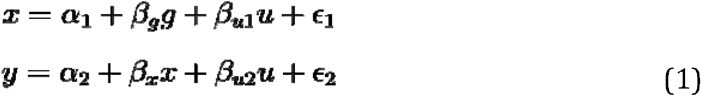

The *α*_⍰_ terms represent intercepts and the _⍰_ terms denote random errors, which are assumed to be independent and normally distributed with mean zero (given g, x and u). This relationship is visualized in the directed acyclic graph (DAG), Figure 1. *x* is a collider on the path from *g* to *y* via *u*, and hence unmeasured confounding between *x* and *y* does not induce spurious associations between *g* and *y* [2]. The advantage of reasoning with DAGs is that the conclusion holds in a nonparametric sense; that is, it holds even when the models are not linear as in equation (1). However, in spite of this, unmeasured confounding may affect the magnitude of a non-zero correlation between g and y, and therefore also the comparison of correlations between g and x versus g and y. This may occur because unmeasured confounding affects the phenotypic variances, and may in particular make the variance of y smaller than the variance of x as seen below. Understanding this artefact of the MR Steiger approach could prove helpful for understanding and utilizing the approach.

**Figure 1:**
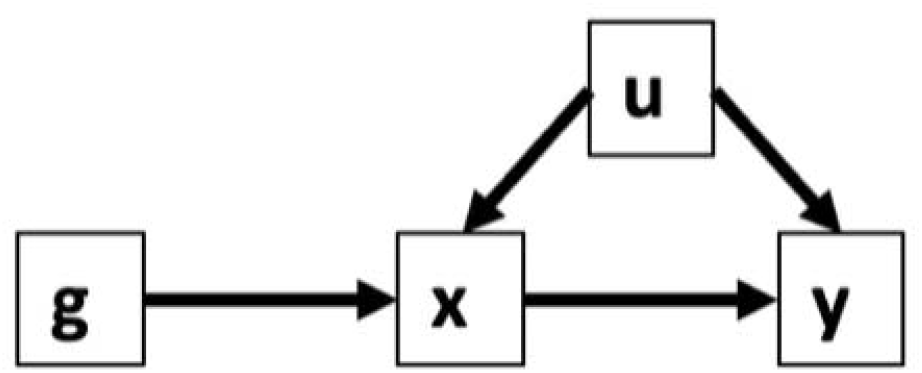
In the DAG below, *g* is the SNP, *x* is phenotype 1, *y* is phenotype 2, and *u* is the unmeasured confounder. Note that *x* is a collider on the path from *g* to *y* via *u*, and hence the association between *g* and *y* is the same regardless of whether there is unmeasured confounding between *x* and *y*.

Assuming the standard regression framework and equation (1), then

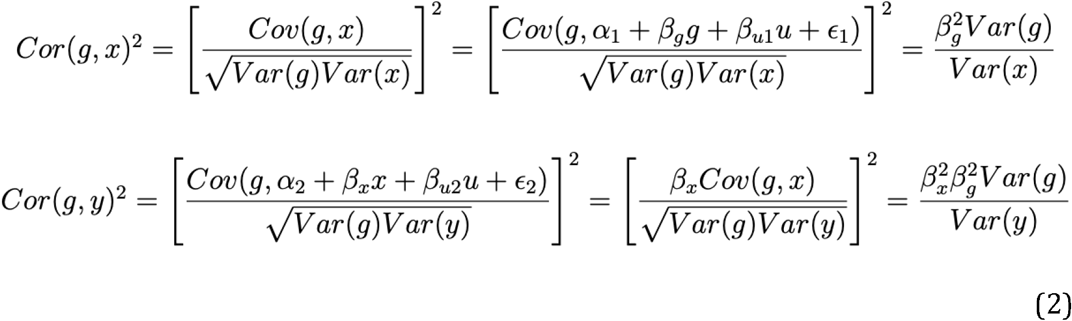

Therefore, the covariance between *g* and *y* does not depend on *β*_*u*2_ and is thus the same, regardless of whether there is unmeasured confounding. The more general reasoning under the DAG in Figure 1 confirms that unmeasured confounding does not induce spurious associations between *g* and *y*. In spite of this, the comparisons of the correlations may be affected by unmeasured confounding. Indeed, using equation (1), *y* can be rewritten such that

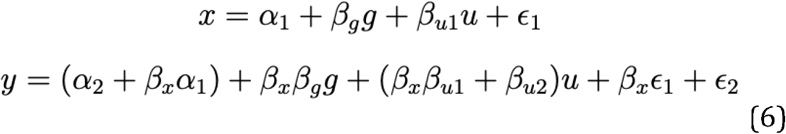

If the signs of *β*_*u*2_ and *β_x_β*_*u*1_ differ (i.e. *β*_*u*2_>0 and *β_x_β*_*u*1_<0 or *β*_*u*2_<0 and *β_x_β*_*u*1_>0), then the unmeasured confounder increases the variance of x but may sometimes decrease the variance of y. In particular, if 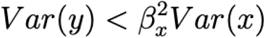 then 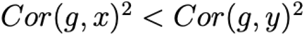 and the false direction may be detected. This occurs when unmeasured confounding biases the effect of x on y towards the null to a sufficient degree. This is an artefact of working with correlations, which do not have the property that they shrink as one moves further in the causal chain.

Consider the following simple simulation study. For n= 100,000 subjects, we generated the SNP *g* and the unmeasured confounder *u* from standard normal distributions. x and y are generated from equation 1 such that the intercepts equal 0 (i.e. *α*_1_ = 0 and *α*_2_ = 0), the genetic effect size, *β_g_*=1, and the effect of *x* on *y, β_x_*=1. The random errors *ϵ*_1_ and *ϵ*_2_ are generated from standard normal distributions. We considered *β*_*u*1_= −5 and vary *β*_*u*2_ from 0 to 12. As seen in Figure 2, there is a range from about 0.1 to 9.5 for *β*_*u*2_ such that estimated *Cor*(*g,x*)<*Cor*(*g, y*) and the MR Steiger approach detects the wrong causal direction. While we considered only one example here, we have created an R package that runs these simulations scenarios for various parameters. The R package called UCMRS is available on GitHub (https://github.com/SharonLutz/UCMRS).

**Figure 2:**
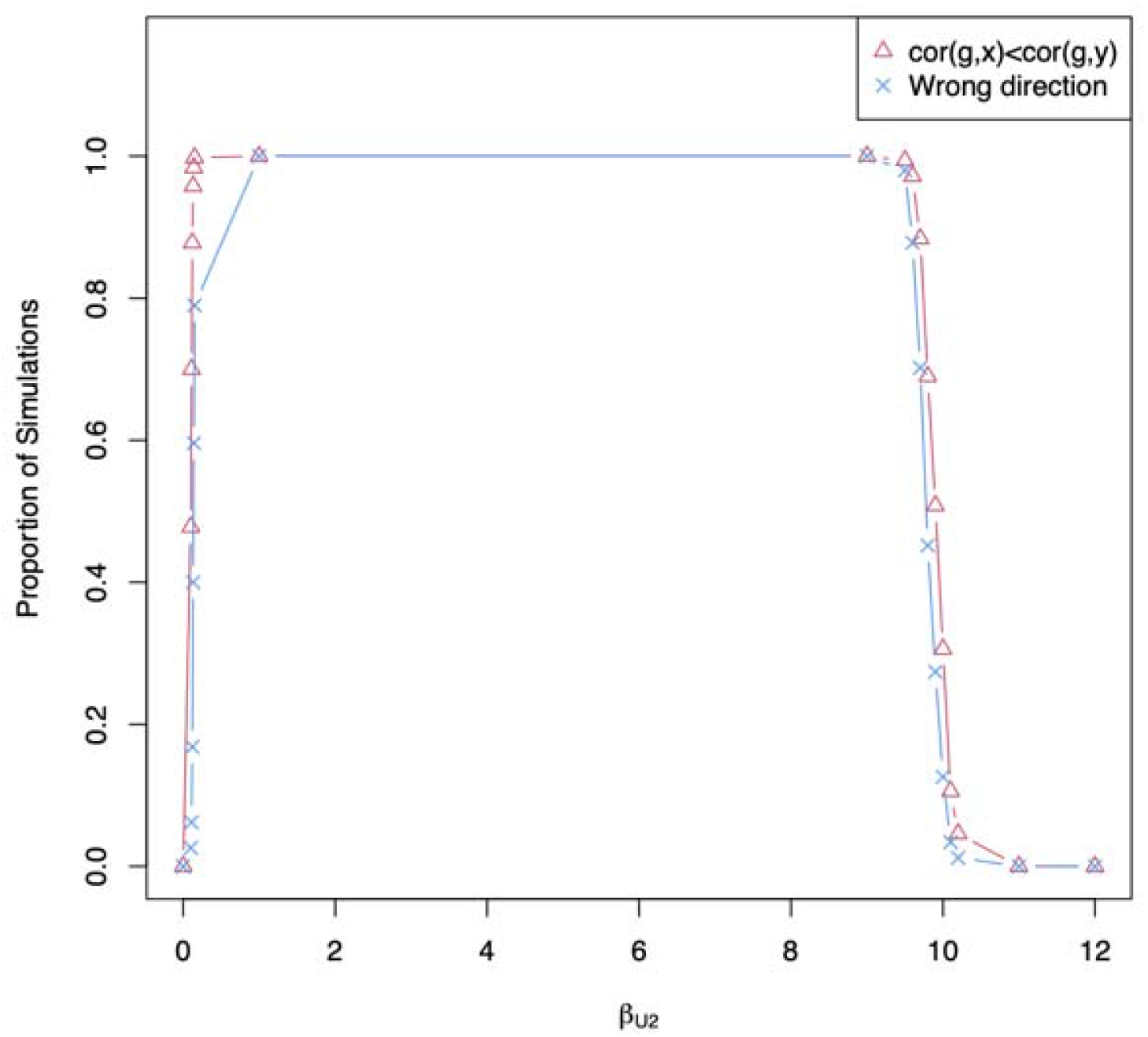
Proportion of simulations where the estimated *Cor*(*g,x*)<*Cor*(*g,y*) and the MR Steiger approach concludes the wrong direction (i.e. y->x).

These results call for caution in using the MR-Steiger method to infer causal directions. It is indeed rarely the case that the association between 2 phenotypes is unconfounded; in fact, concerns about confounding form the key motivation behind MR methods. In line with the original MR Steiger manuscript [1], caution is especially warranted when unmeasured confounding is expected to induce a bias in a direction opposite to the effect between both phenotypes. Other major concerns about the MR-Steiger method come from the assumption that *g* has no direct effect on one of the phenotypes. [1, 2] Indeed, in settings where it is not a priori known whether x influences *y* or *y* influences *x*, there can be no biological basis to conclude that *g* has no direct effect on either *x* or *y*, since assumptions about the absence of a direct effect demand knowledge as to whether *x* influences *y* or vice versa. This makes the MR-Steiger method reliant on strong mathematical convenience assumptions. Sensitivity analyses are needed to study the effects of confounding by unmeasured variables and pleiotropy on the MR Steiger approach.

## Acknowledgements

Research reported in this publication was supported by the National Institutes of Health grants K01HL125858 (SML), NICHD R01HD085993 (ACW), and the Cure Alzheimer’s Fund (CL). We would like to thank Dr. Kate Tilling for helpful discussions and for suggesting the considered simulation study example.

